# A versatile CRISPR-based system for lineage tracing in living plants

**DOI:** 10.1101/2023.02.09.527713

**Authors:** Mattia Donà, Gabriele Bradamante, Zorana Bogojevic, Ruben Gutzat, Susanna Streubel, Magdalena Mosiolek, Liam Dolan, Ortrun Mittelsten Scheid

## Abstract

Individual cells give rise to diverse cell lineages during the development of multicellular organisms. Understanding the contribution of these lineages to mature organisms is a central question of developmental biology. Several techniques to document cell lineages have been used, from marking single cells with mutations that express a visible marker to generating molecular bar codes by CRISPR-induced mutations and subsequent single-cell analysis. Here, we exploit the mutagenic activity of CRISPR to allow lineage tracing within living plants. Cas9-induced mutations are directed to correct a frameshift mutation that restores expression of a nuclear fluorescent protein, labelling the initial cell and all progenitor cells with a strong signal without modifying other phenotypes of the plants. Spatial and temporal control of Cas9 activity can be achieved using tissue-specific and/or inducible promoters. We provide proof of principle for the function of lineage tracing in two model plants. The conserved features of the components and the versatile cloning system, allowing for easy exchange of promoters, are expected to make the system widely applicable.

**SIGNIFICANCE STATEMENT:** By targeting Cas9 in a tissue- and time-specific way to correct a frameshift mutation, resulting in fluorescence labelling of nuclei, we generated a method for *in vivo* visual lineage tracing in two model plants. The versatile cloning system makes the system widely applicable in other plants.

## INTRODUCTION

Although attempts are being made to create artificial cell models, Rudolf Virchow’s statement (1855) “*omnis cellula e cellula*” (all cells originate from another cell) is still valid. It refers to the connection in space and time between cells within an organism. How cell lineages coordinate to generate multicellular organs and organisms is a central question of developmental biology. Following cell-to-cell continuity from zygotes to early embryonal stages and tracking cell lineages in later stages of multicellular organisms can be a difficult and complex challenge. Documenting a complete lineage map of an adult animal was a breakthrough but remained an exception for *C. elegans*, available only due to the limited and definite cell number and a fast and stringent developmental program in this nematode (Sulston *et al.* 1983). In organisms with billions of cells that form complex tissues, grow slowly, have mobile cells, or plastic development, cell lineage tracing requires labeling precursor cells and ways to follow the label in all derivative cells. This has been achieved in many organisms by numerous techniques, with different advantages, but none of them free from limitations (reviewed in Buckingham and Meilhac 2011, Garcia-Marques *et al.* 2021).

Most widely applied are optical methods combining reporter genes with genetic components that change their expression in a time-, tissue- or trigger-dependent manner. Popular reporter genes encode enzymes like lacZ or GUS/uidA that convert substrates into coloured products, detectable after histological staining. Their benefits are cell-autonomous expression and visibility of the signal even upon low expression of the reporter. Still, they require tissue fixation, which is incompatible with observation in living organisms. This limit is overcome by using fluorescent proteins (FPs), with manifold advantages (Yagi *et al.* 2021). FPs are encoded by relatively short genes, making transgene design easier, and they can be fused as tags to other proteins, often without functional interference. They represent a growing family of proteins with different excitation and emission wavelengths, have different signal strengths and half-lives, and can be targeted to different compartments of the cell. In the context of lineage tracing, their fusion to histones is especially convenient, as this guarantees high amounts of FP-tagged proteins resulting in strong signals, containment in the nucleus, and faithful propagation of the label to each daughter cell. FPs can be recorded in living tissues, with time lapse observation (e.g., Reddy *et al.* 2004) or live imaging over longer periods (e.g., von Wangenheim *et al.* 2016).

FPs are often combined with genetic components that switch the activity of the reporter genes in a controlled way. These are endonucleases like recombinases and, in a growing number of cases, CRISPR-based gene editors. In both cases, targeted modifications at the reporter genes are controlled by transcribing the endonuclease genes from cell- or tissue-specific promoters or after chemical or physical induction. The toolbox contains simple off/on switches of reporters, FP colour switches, but also systems to consecutively label cells generations or accumulate spontaneous or endonuclease-induced mutations. The latter approaches have created the chance to complement optical lineage tracing with single-cell sequencing approaches, even in complex animal organs (reviewed in Garcia-Marques *et al.* 2021).

Several techniques for lineage tracing that are successful in animals are not or not easily applicable in plants (Buckingham and Meilhac 2011). However, plants have some features that support investigating cell lineages (Poethig 1989). One advantage is that, with rare exceptions, plant cells are immobile. New cells stay connected with their mother cells by common cell walls, allowing very reliable lineage tracing at high resolution. In contrast to most animals, organ formation in plants continue throughout lifetime with high plasticity. While this can lead to variable cell lineages, it allows studying fundamental developmental decisions in response to environmental stimuli.

Early studies on cell lineage in plants took advantage of chlorophyll as a natural pigment in many tissues, allowing a non-invasive, non-transgenic approach. The presence or absence of chloroplasts, loss of genes required for chlorophyll biosynthesis, or defects in nuclear-plastid interaction distinguish a white cell and all its progeny from a green background and enable sector analysis with a natural visible marker (Furner and Pumfrey 1992, Irish and Sussex 1992, Kato *et al.* 2007, Pogorelko *et al.* 2016, Rédei 1963). However, unbiased observation of cell lineages requires that the presence or absence of the label does not interfere with regular development. This is a limit for chlorophyll variegation, as it likely has drastic effects on the metabolic state of the cells. Other, less essential natural pigments have been utilized repeatedly in connection with mobile genetic elements. Insertion or excision of transposons in genes that determine pigmentation represents random events in somatic cells that mark this progenitor and all its progeny cells (Becraft 2013). Colour sectors contributed to the discovery of mobile elements (McClintock 1951) and were later frequently used to study development (e.g., Hernandez *et al.* 1999, Inagaki *et al.* 1994). Refining the lineage tracing by controlling the mobility at defined developmental stages became possible by including transposon-derived elements in transgenic reporter constructs. Their excision can be induced by trans-activation, e.g., by heat activation of the autonomous Ac leading to excision of a non-autonomous Ds element between a promoter and a reporter gene or similar combinations (e.g., Barkoulas *et al.* 2008, Kurup *et al.* 2005, Torres-Martínez *et al.* 2020). Controlled activation of recombinases can excise or flip matching target sequences in reporter genes and thereby label a cell lineage (e.g., Efroni *et al.* 2015, Shi *et al.* 2019, Suzuki *et al.* 2020), or eliminate lineages, ablating precursor cells by expressing diphtheria toxin (Glowa *et al.* 2021). Targeting a key gene in pigment biosynthesis with CRISPR-Cas9 created sharp sectors lacking colour in the petals of Japanese morning glory and allowed to study periclinal chimaera (Watanabe *et al.* 2017). In case that stable transformation of reporter genes and endonucleases is not (yet) established for the plant under investigation, irradiation to induce visible sectors (Harrison *et al.* 2007), or transient transformation of single cells with reporter constructs (Bossinger and Spokevicius 2018) have been successfully applied. Different reporter systems based on stably integrated transgenes are established in many model plants (e.g., Schürholz *et al.* 2018, Smetana *et al.* 2019, Van Ex *et al.* 2009, Wang *et al.* 2020), and single-cell analyses are feasible and become popular also in plants (reviewed in McFaline-Figueroa *et al.* 2020). However, the advantage of the cell wall-determined spatial fixation of plant lineages is lost in single cell approaches as these require protoplast preparation or nuclei isolation. Therefore, to reflect dynamic processes over time, optical tracking in the context of living plant tissue is the method of choice and should combine strong, stable fluorescence signals visible after reporter gene switches that are controlled in time, location, and frequency and lead to non-reversible changes present in all progeny cells.

Here, we describe a versatile system in which the mutagenic activity of Cas9 is used to allow lineage tracing in living material, in different tissue and organs, and at different developmental stages. It is based on the repair of a Cas9-induced double strand break by non-homologous end joining, resulting in a gain of function of a fluorescent protein that is stably integrated into nuclei. This provides a strong signal in the initial cell and is propagated during all subsequent cell divisions. By controlling the endonuclease activity of Cas9 with cell- or tissue type-specific promoters or chemical induction, the starting point of the lineage tracing can be chosen variably and precisely. We provide proof of principles for long-term lineage tracing in the model plants *Arabidopsis thaliana and Marchantia polymorpha*. The high conservation of histone as a reporter protein and the versatile cloning system will render the principle widely applicable across plants and likely other eukaryote kingdoms.

## RESULTS

### Design principles of the reporter lines

Lineage tracing in living tissue over longer time periods requires observation by non-destructive imaging of strong and stably inherited signals. We have constructed a reporter system to label the nuclei of individual cells and all their descendants, based on correcting a frameshift after a CRISPR/Cas9-induced double strand break. The readout is nuclear-localized fluorescence provided by mClover fused to the C-terminus of histone H2B, an abundant and strongly conserved nucleosome subunit. Early after the start codon of the ORF, we inserted a part of a sequence from the human gene *AAVS1* containing a well-characterized CRISPR/Cas9 target site (Mali *et al.* 2013) with no homology in the Arabidopsis reference genome (Figure 1a). We generated three different reporter constructs. In reporter LT0, we added 21 bp of the *AAVS1* gene sequence maintaining the reading frame with H2B-mClover (LT0). Cells expressing this construct are expected to show stable nuclear fluorescence. In LT1 or LT2, we added either 20 bp or 22 bp of the *AAVS1* gene sequence respectively, creating frameshifts of either +2 or +1 within the ORF so that premature stop codons prevent translation of functional H2B-mClover. Cells expressing either construct are therefore expected to have no nuclear fluorescence, unless a frameshift mutation occurs. The LT reporters are controlled by the promoter of the HTR5 gene, providing high, cell cycle-independent, and ubiquitous expression, except in the central cell of the female gametophyte (Ingouff *et al.* 2017). The LT reporter cassettes are schematically shown in Figure 1a, the binary vectors used to generate plants with different reporter systems in Figure 1b and c. All vectors were constructed with the modular GreenGate cloning system (Lampropoulos *et al.* 2013), which allows easy exchange of the elements and makes the vectors versatile and adaptable for different purposes. We combined the double strand break-inducing spCas9 variant with three different promoters well characterized in *Arabidopsis thaliana*: pHTR5 (described above) for constitutive expression (pHTR5Cas-LT0/1/2), pGL2 for expression in the atrichoblast lineage of the root epidermis (Di Cristina *et al.* 1996, Masucci *et al.* 1996) in the reporters pGL2Cas-LT0/1/2, or pCLV3 for expression in the stem cells of the shoot apical meristem (SAM) (Laux 2003) in the reporters pCLV3Cas-LT0/1/2. All vectors contain a cassette expressing a gRNA targeting the *AAVS1* insert under the control of the U6-26 snRNA promoter, designed to allow multiplex CRISPR/Cas9 cleavage (Xie *et al.* 2015). Cleavage in the *AAVS1* region occurs only when Cas9 and gRNA are expressed together. Subsequent repair by the error-prone non-homologous end joining pathway very frequently creates small indels (Weiss *et al.* 2022, Zhang *et al.* 2020) that can result in the correction of the frameshifted H2B-mClover reading frame. Nuclei of cells with such events will be labelled by mClover fluorescence. As the mutation will modify the gRNA target sequence, the site is protected from further cuts. Should the cell divide, all descendants will inherit the restored functional frame and have labelled nuclei, visualizing lineages within the tissue context of living plants also when Cas9 is no longer expressed. We constructed additional vectors bearing only the gRNA and LT reporter cassettes (LT0/1/2), as controls for measuring the background frequency of spontaneous frameshift correction in the absence of Cas9 and for sequential transformation with the constructs described below. In another set of constructs, we added a module that labels the plasmalemma with mRuby-SYP132, a ubiquitously expressed membrane trafficking protein (Enami *et al.* 2009) (Figure 1b). We also generated constructs that allow the expression of cell type-specific Cas9 only after induction by external application of estradiol, using one of the three promoters to drive the XVE transcription factor binding to the LexA site cloned upstream of *spCas9* (ipHTR5Cas, ipGL2Cas, ipCLV3Cas, Figure 1b). For analysis of lineages in the SAM, we combined the LT reporters with another marker in which the CLV3 promoter visualizes the stem cell domain in red (pCLV3:H2B-mCherry) (Gutzat *et al.* 2020). All constructs have a cassette allowing visual selection of successful transgenesis by seed fluorescence, and all elements are between left and right border sequences for T-DNA integration (Figure 1b). To investigate the suitability beyond *Arabidopsis thaliana*, we adapted the constructs to render them suitable for lineage tracing in *Marchantia polymorpha*, as the meristematic stem cell regulation has similarities, but also differences with that of Arabidopsis (Hata and Kyozuka 2021). The promoter of the *MpEF1a* gene was inserted upstream of the H2B-mClover sequence for strong and constitutive expression (Althoff *et al.* 2014) of the reporter cassettes. We used pMpEF1a also to drive Cas9 in constructs pEF1aCas-LT0/1/2, or MpYUC2 that provides expression in meristematic cells (Eklund *et al.* 2015) for construct pYUC2Cas-LT0/1/2, together with a selection cassette that confers resistance to hygromycin to transgenic lines (Figure 1c). The Marchantia construct were also modified to control cell type-specific Cas9 after estradiol application, resulting in plasmids ipEF1aCas-LT0/1/2 and ipYUC2Cas-LT1/1/2 (Figure 1c).

**Figure 1.**
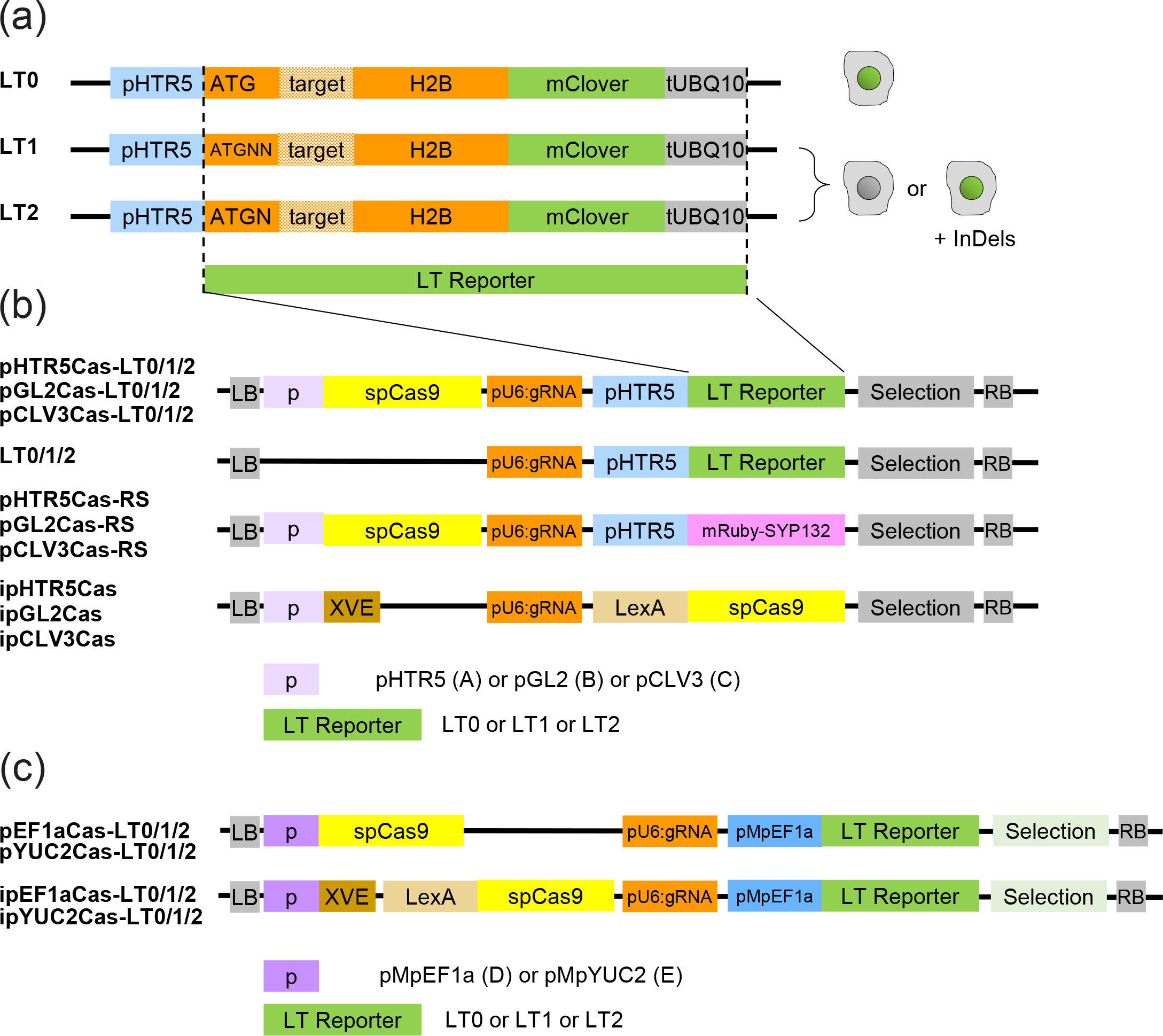
Schematic representation of constructs for the lineage tracing system. (a) Composition of the reporter cassette. The lineage tracing reporters contain the protein-coding sequence of histone 2B (H2B) fused to the fluorescence protein mClover, under the control of the Arabidopsis HTR5 promoter for strong and constitutive expression and the terminator of the *UBIQUITIN* 10 gene (UBQ10t). A part of the human gene *AAVS1*, not present in the Arabidopsis genome, is inserted as gRNA target sequence soon after the start codon ATG, either in frame with H2B-mClover (LT0), or with 2 (LT1) or 1 (LT2) additional bps, respectively. LT0 is expected to provide constitutive labelling in all nuclei, while the frameshifts in LT1 and LT2 prevent H2B-mClover translation. If LT1 or LT2 are combined with CRISRP/Cas9 and gRNA targeting the AAVS1 sequence, indels resulting from repair by non-homologous end joining can restore the frameshift, e.g., by indels of +1 bp or −2 bp for LT1, or +2 bp or −1 or −4 bp for LT2. After the error-prone repair, the gRNA target sequence is resistant to subsequent cleavage, and the translatable LT module is inherited by all descendants of the cell with the original frameshift correction. (b) Vectors constructed with the GreenGate modular assembly for lineage tracing in Arabidopsis thaliana. Between the left and right border (LB, RB) for T-DNA integration, they contain a cassette for gRNA transcription from the U6-26 promoter (pU6:gRNA) and a cassette for selection of transgenic plants, conferring fluorescence to seeds. The double strand break-producing endonuclease spCas9 is under control of different promoters (p), namely pHTR5 for constitutive expression, pCLV3 for expression in the stem cells of the shoot apical meristem, or pGL2 for expression in the atrichoblast lineage of the root epidermis. The vectors pHTR5Cas-LT1/2, pGL2Cas-LT1/2, or pCLV3Cas-LT1/2 allow recording frame shift correction events in the whole plant, in the root epidermis, or the stem cells of the shoot apical meristem, respectively. Vectors LT0/1/2 lack Cas9. LT0 confirms constitutive expression of the reporter independent of Cas9; LT1/2 measure the frequency of background events restoring the frameshift. The -RS vectors provide constitutive or tissue-specific Cas9 expression together with constitutive expression of mRuby-SYP132, labelling the plasma membrane; they are applied in combination with LT1/2-expressing plants. In vectors ipHTR5Cas, ipGL2Cas, or ipCLV3Cas, Cas9 expression depends on induction by estradiol, as it will be only produced if the XVE transcription factor binds to the LexA site. Combined with LT1/2, correction of the frameshift can be programmed in specific cells at specific time points. (c) Vectors constructed with the GreenGate modular assembly for lineage tracing in *Marchantia polymorpha*. Between the left and right border (LB, RB) for T-DNA integration, they contain the cassette for gRNA transcription from the U6 promoter (pU6:gRNA), LT reporter module LT1 or LT2 fused to pMpEF1a for strong constitutive expression as well as a cassette for selection of transgenic Marchantia plants conferring resistance to hygromycin. The vectors pEF1aCas-LT1/2 and pYUC2Cas-LT1/2 contain the SpCas9 gene under the control of either pMpEF1a (as described above) or MpYUC2 for expression in the meristemoid. Vectors ipEF1a-LT1/2 and ipYUC2Cas9-LT1/2 allow Cas9 expression upon induction with estradiol as described in (b).

### Functional test of the reporter lines

Constitutive expression of the reporter lines in the tissue of interest is a prerequisite for the system to work. This was confirmed in plants containing the reporter with a functional ORF for the H2B-mClover but missing the Cas9 cassette (LT0) by strong nuclear fluorescence in all cells in leaves, roots, and meristems (Figure 2a). The fluorescence signal was also stably inherited in the second and third generation of the transgenic lines, excluding silencing. The plants with the reporters are phenotypically indistinguishable from the wild type (Figure 2b).

**Figure 2.**
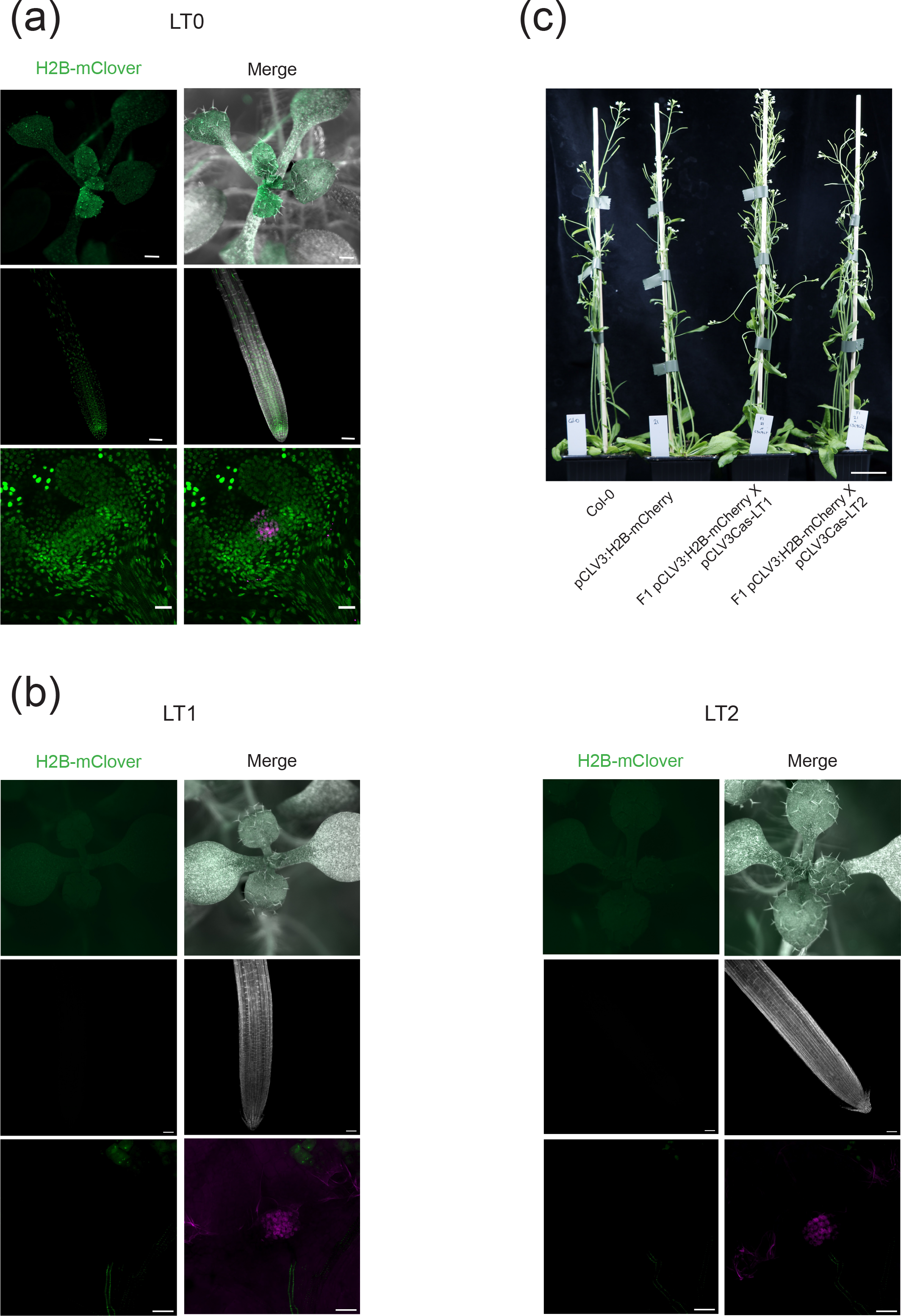
Reporter expression stability and background control. (a) Representative images of Arabidopsis plants with reporter LT0 at 12 days after germination (dag), whole seedlings (top), roots (middle), or shoot apical meristems (SAMs) (bottom). (b) Representative images of Arabidopsis plants with reporters LT1/2 but lacking Cas9 expression at 7 dag. Left: H2B-mClover (green); right: Merge, green channel combined with visible light images and magenta channel to detect plasma membrane staining (root) or the stem cells (SAM). (c) Phenotype of plants transgenic for the lineage tracing reporter constructs. Bars = 500 μm (a and b) or 3 cm (c).

Another prerequisite for the functionality of the system is the strict dependence of fluorescence signals on the presence of the Cas9 endonuclease. Plants containing the reporters LT1 or LT2 with the frameshift but lacking the Cas9 cassette (LT1 or LT2) do not contain any labelled nuclei, excluding a background of spontaneous frameshift restoration (Figure 2c). By contrast, the combination of Cas9 expression and reporter LT1 or LT2 results in labelled nuclei, seen as either individual fluorescent nuclei directly after the restoration, as row of labelled cells after a few subsequent divisions, or as large fluorescent sectors at later stages (Figure 3a-f).

**Figure 3.**
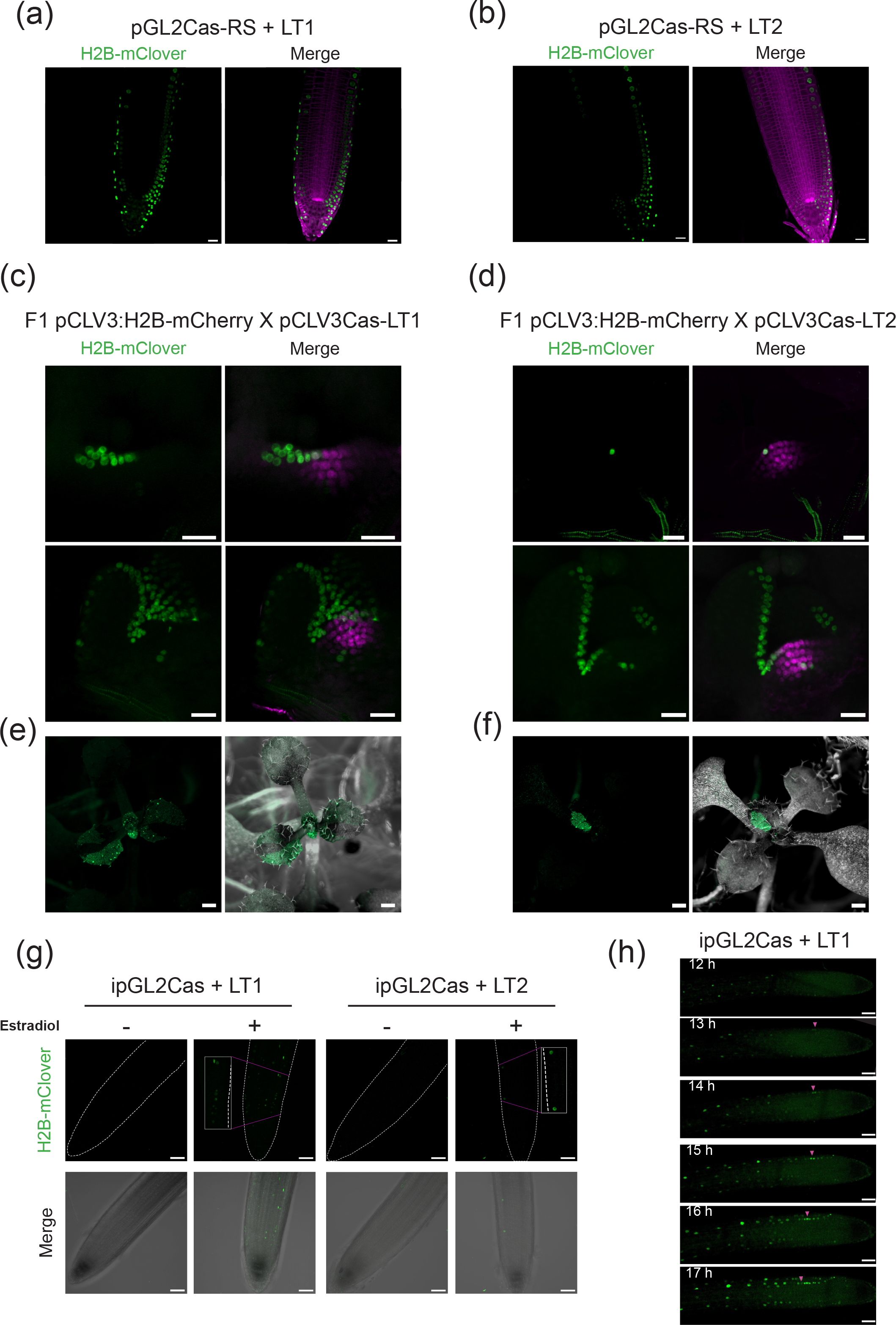
Functional test of the reporter lines. Representative images after initial frameshift corrections and single lineages formed later in Arabidopsis. (a, b) Roots 7 dag expressing pGL2Cas-RS introgressed in plants expressing LT1 or LT2 (b) reporters. (c, d) Maximum intensity projections of SAMs from 7 day-old F1 hybrids between stem cell reporter line pCLV3:H2B-mCherry (magenta) and lineage tracing reporter lines pCLV3Cas-LT1 (c) or -LT2 (d) (green). Top: soon after frameshift correction (a single nucleus in (d)); bottom: lineages of labelled nuclei originating after cell division. (e, f) Sectors with labelled nuclei in aerial tissue of 12 days-old seedlings. (g) Images of the root epidermis from plants with the inducible Cas9 vectors after preceding incubation with or without estradiol. (a-g) H2B-mClover and Merge as in Figure 2. (h) Time-lapse imaging of an Arabidopsis root 10 dag bearing ipGL2cas and LT1. Arrows (magenta) mark nucleus with initial frameshift correction and subsequent formation of the lineage. Bars = 50 μm (a, b, g, h), or 20 μm (c, d) or 500 μm (e, f).

To support the visualization of the tissue context, we additionally labelled the plasma membrane by expressing Ruby-SYP132 (mentioned above) together with the LT1 or LT2 reporter (Figure 3a, b). We applied this when we tested the construct variants where Cas9 expression depends on the external application of estradiol. To validate our construct design and the optimal condition for Cas9 induction, we used lines bearing ipGL2-Cas, as it provides an easy and reliable readout in roots. While no fluorescent nuclei are observed without induction, an increasing number of events are seen during several hours after estradiol application (Figure 3g). Live imaging of the same root after 2 hours estradiol induction illustrates the progression of labelling and allows following the lineage along a timeline with first labelled nuclei visible 11 hours after the treatment (Figure 3h).

Taken together, these data demonstrate that the reporter system is suitable, reliable, and versatile to trace lineages derived from single cells during development.

### Recapitulation of known lineages in the root epidermis

To verify that the system faithfully recapitulates well-described cell lineages in plants, we applied it to the root epidermis of *Arabidopsis thaliana* that contains two well-established cell type lineages: the alternating files of root hair cells (H) originating from trichoblasts, and non-hair cells (N) originating from atrichoblasts. Their specification is determined by contact with either one or two neighbouring cortical cells and antagonistic transcription factors and repressors (reviewed in Dolan and Costa 2001). The gene *GLABRA2 (GL2*) encodes a homeobox domain protein that suppresses root hair formation in the atrichoblasts (Di Cristina *et al.* 1996) and is active during the early differentiation of cells in the root meristem (Dolan *et al.* 1998). We used the promoter of *GL2* to drive the Cas9 expression in combination with the LT1 or LT2 reporter and a mRuby-SYP132 fusion to label plasma membranes. Indeed, we observed labelling in adjacent cell files corresponding to atrichoblasts in the root epidermis (Figure 4a-d). In some cases, these cell files originate close to epidermal initials at the root apex (Figure 4b). These results are in accordance with previously shown patterns of GUS reporters for *pGL2* and single-cell root transcriptomics data that document *GL2* expression early during epidermal lineages differentiation (Masucci *et al.* 1996). We also generated a new translational reporter for direct comparison of *GL2* promoter activity, by expressing an H2A-BFP under the control of pGL2 (Figure 4e). Its pattern overlaps with the pGL2 lineage tracing signals, but with weaker fluorescence. Therefore, we demonstrate that the CRISPR-based lineage tracing system can be used to define lineages from early to late stages of differentiation and development, resulting in strong nuclear labelling only in cells where the driving promoter has been shown to be active.

**Figure 4.**
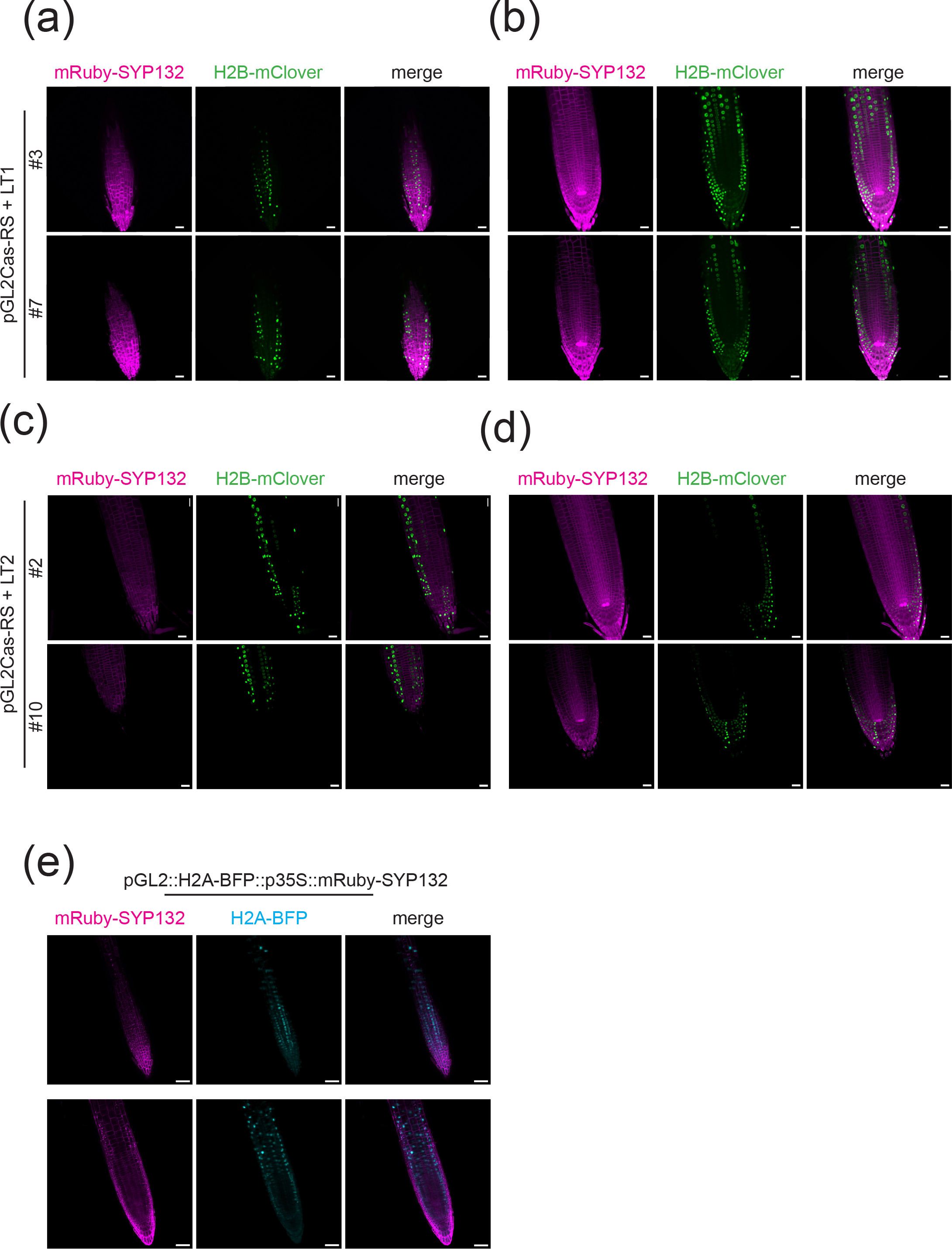
Lineage tracing in the root epidermis of Arabidopsis. Representative images of roots in seedlings 4 dag bearing pGL2Cas-RS together with LT1 (a, b) or LT2 (c, d), respectively. Maximum intensity projections of the root epidermis (a, c) or 10 μm sections in the middle of the root (b, d). # = individual transgenic lines. (e) Transcriptional reporter for GL2 expression via localization of H2A-BFP (blue) under the control of the GL2 promoter, together with expression of plasma membrane marker mRuby-SYP132 (magenta). Merge (right): combined images of plasma membrane staining with mRuby-SYP132 (magenta, left) and H2B-mClover (green) or H2A-BFP (blue). Bars = 10 μm.

### Visualization of lineages in the shoot apical meristem

To test the applicability of the lineage tracing reporters for less known cell lineages, we modified the system to label stem cells in the shoot apical meristem (SAM). This tissue forms all above-ground postembryonic plant organs, including flowers. Therefore, at least a subset of these stem cells is part of a functional germline (reviewed in Burian 2021, Nguyen and Gutzat 2022). The structure of the SAM and regulatory mechanisms determining the size of the stem cell pool are well characterized. The separation of the SAM into three distinct layers (L1, L2, L3) with specific orientation of dividing cells and different frequencies of karyotype mutations is known for long (e.g., Satina *et al.* 1940). However, the stem cells as defined by expression of the signalling peptide CLAVATA 3 (CLV3) span all three layers in the Arabidopsis shoot meristem (reviewed in Nguyen and Gutzat 2022). Therefore, we used the promoter of *CLV3* to express Cas9 in combination with LT1 or LT2. To increase the spatial resolution of frameshift corrections occurring in the SAM, Arabidopsis plants were grown on soil under short-day conditions for 21 days. These conditions let plants develop a larger apical meristem (Burian *et al.* 2022). Images of apices showed labelling of cell files emerging from different layers of the SAM and expanding into diverse aerial tissues of Arabidopsis (Figure 5a, b). The frequency of labelling events with the LT1 reporter is higher than for LT2. At later developmental stages, the H2B-mClover labelled nuclei originating from single events in the CLV3 domain allow tracking the division and differentiation history of specific SAM lineages (see also Figure 3a). This demonstrates that the CRISPR-based reporters are suitable to trace the origin and developmental branches in the SAM, and potentially in every cell type where differentiation determination takes place.

**Figure 5.**
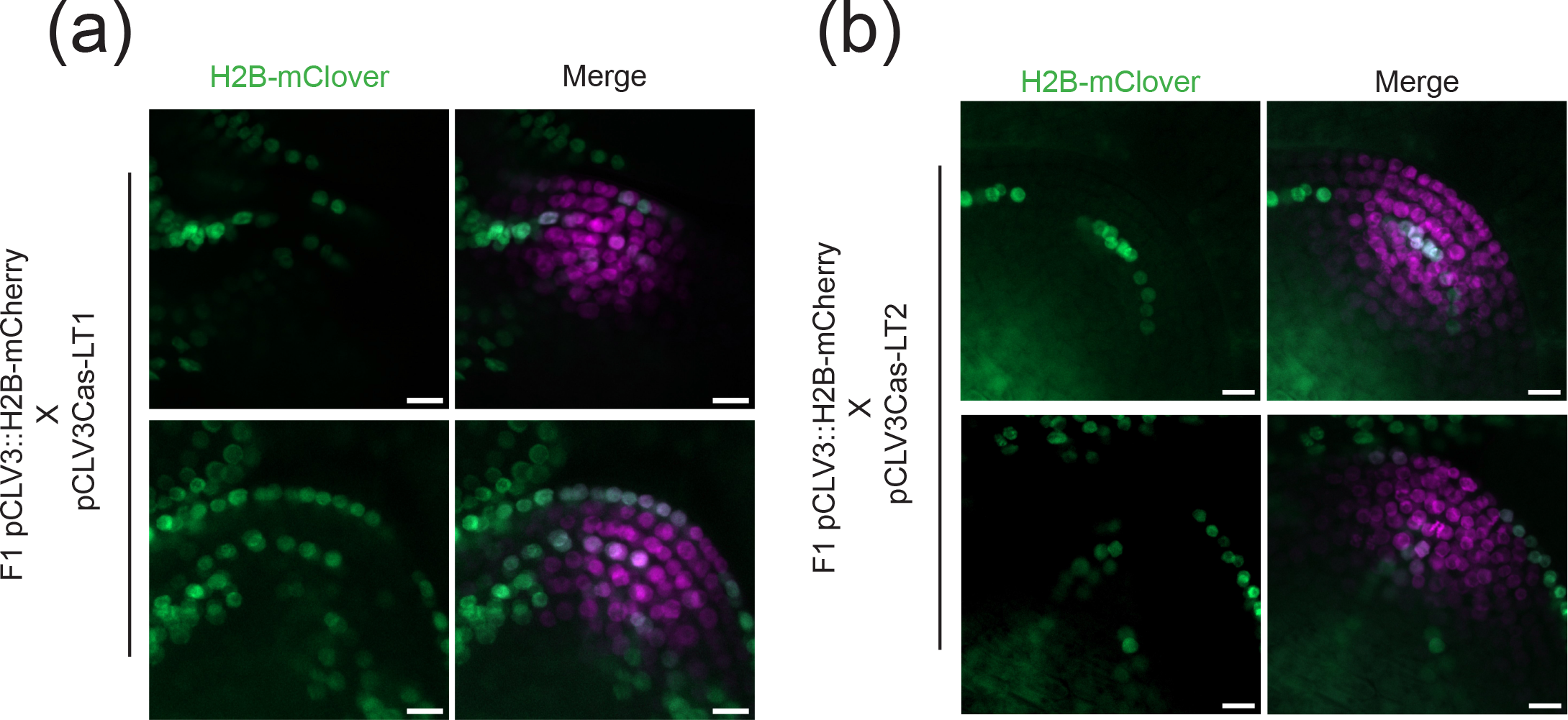
Lineage tracing in the shoot apical meristem of Arabidopsis. Maximum intensity orthogonal projection of shoot apical meristems from plants 21 dag grown under short day conditions. (a, b) Examples of SAMs from F1 hybrid seedlings carrying the CLV3 domain marker pCLV3∷H2B-mCherry (magenta) and the pCLV3Cas-LT1 (a) or the pCLV3Cas-LT2 (b) reporter (green). Bar = 10 μm.

### Frequency of events and maintenance of the reporter lines

Once the CRISPR/Cas9-induced double strand break in the AAVS1 target is repaired and mutated, regardless of whether correcting the frameshift or not, the reporters are not responsive anymore. Although the frequency of frameshift corrections is high enough to see events in nearly all pCLV3Cas-LT1/2 plants, the transmission frequency of a functional reporter to the next generation was high, suggesting that expressing the Cas9 protein fromthe CLV3 promoter yields relatively little germline events of reporter activation. To quantify the frequency of reporter activation in the germline, we crossed the LT reporter lines with the stem cell reporter line, resulting in an F1 progeny with a single copy of the LT target gene. We then scored 52 F1 plants from six independent crossings of each reporter line for visible fluorescence sectors in the early stage, resulting in 69% and 71% positive seedlings for LT1 and LT2, respectively, which had inherited a functional reporter without mutations in the germline (Figure 6a). Presence of fluorescent nuclei in the whole seedlings including embryonic tissue, indicating that the reporter had been activated in the germline of the parent plant, were found with 31% and 29%, respectively. Next, we grew F1 plants to maturity and scored seedlings from 10-37 independent F2 populations for fully fluorescent seedlings, seedlings with developing sectors, or seedlings without any fluorescence (Figure 6b). We scored 10.1% and 3.4% frameshift-restoring germline events for LT1 and LT2, respectively. Pools without any fluorescence, indicating germline mutations not correcting the frameshift, occurred with ratios of 3.9% and 16.7% for LT1 and LT2, respectively. Therefore, plants that had lost functional reporters either way represent a small portion of the progeny, can be easily identified at the early seedling stage and discarded during propagation of the lineage tracing lines.

**Figure 6.**
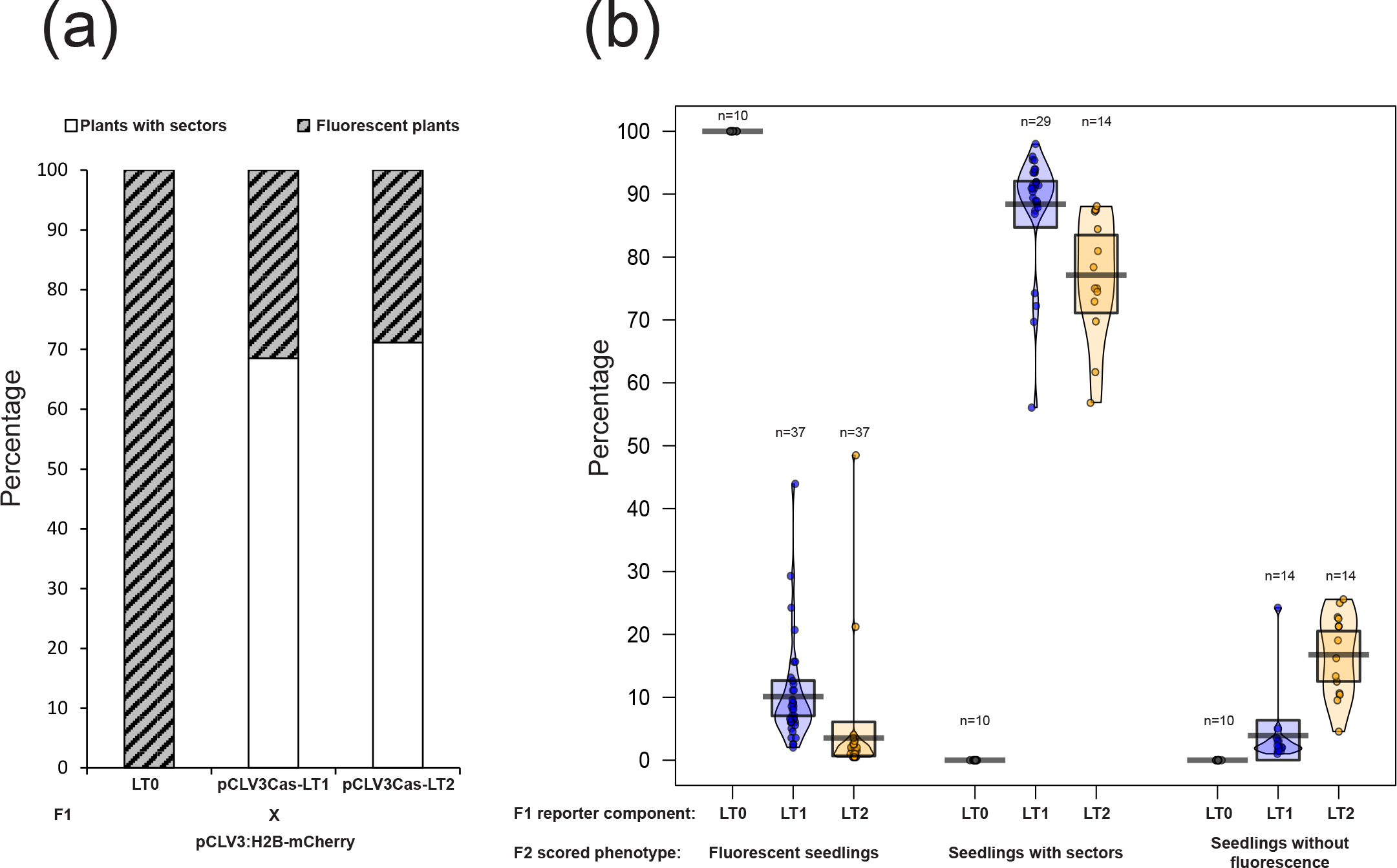
Maintenance of functional reporters across generations. Percentage of plants carrying a functional reporter (plants with sectors) in the F1 and F2 generations. (a) Percentage of F1 plants (n = 57) showing fully labelled nuclei. (b) Percentage of F2 seedlings (n = 200 for each population) obtained from n individual F1 plants showing fully fluorescent seedlings (frameshift correction in the germline), seedlings with sectors (functional reporter maintained), or seedlings without fluorescence (germline mutation without frameshift correction).

### Application in a different model plant, *Marchantia polymorpha*

With the wealth of molecular and genetic data and tools, its suitability for microscopic observation, short generation times, and the ease of transgenesis, *Arabidopsis thaliana* still serves as a major model for developmental studies in plant biology. However, as described above, the presence of two alleles of the reporter gene in inbred lines during most of the life cycle can complicate the lineage tracing. In addition, it is of interest to see developmental processes in the context of evolutionary biology. *Marchantia polymorpha* (Marchantia) is a member of the monophyletic bryophytes. The dominant phase of the life cycle of bryophytes is haploid (for review see Bowman 2022). Like in higher plants, new tissues and organs in Marchantia originate from stem cells in meristems (Hirakawa *et al.* 2020). The knowledge regarding the development of this particular basal plant lineage is limited. Nevertheless, its appeal as a model organism has been strengthened by its reduced genetic redundancy and the conservation of developmentally relevant key genes.Therefore, we asked if the lineage tracing system could be applied also to this species. We exchanged the promoter upstream of the LT0, LT1, orLT2 reporter for two Marchantia-specific promoters: *pMpEF1a* as a strong and constitutive promoter (Althoff *et al.* 2014), and *pMpYUC2*, which is active in the meristematic cells present in the apical notch (Eklund *et al.* 2018). Expression of Cas9 under the control of the *MpEF1a* promoter in the in-frame reporter LT0 results in strong and constitutive labelling of all nuclei in young gemmae (Figure 7a). This demonstrates that the fluorescent signal is stable over the course of plant development (Supplemental Figure 1a). Lines carrying the out-of-frame reporters pYUC2Cas-LT1 and pYUC2Cas-LT2 form sectors with fluorescent nuclei (Figure 7b, c). The sectors originate in the meristems at the notches of the gemmae and are visible as soon as three days after transferring new gemmae on fresh growth medium (Figure 7b, c). Reading frame reconstitution in regions proximal to the meristems, where most of the cell divisions occur, result in fluorescent sectors with a high density of nuclei. Sectors originating more distally, where tissue differentiation dominates, have much fewer fluorescent nuclei in large cells (Supplemental Figure 1b, c). Therefore, the reporters provide information about the time of the mutation in *M. polymorpha* apical notches during division versus differentiation (Hirakawa *et al.* 2020), with a simple visual readout. In summary, these results confirm that the lineage tracing system is functional and easily adaptable to new plant species and likely any organism accessible by transgenesis, providing a valuable platform for addressing manifold developmental questions.

**Figure 7.**
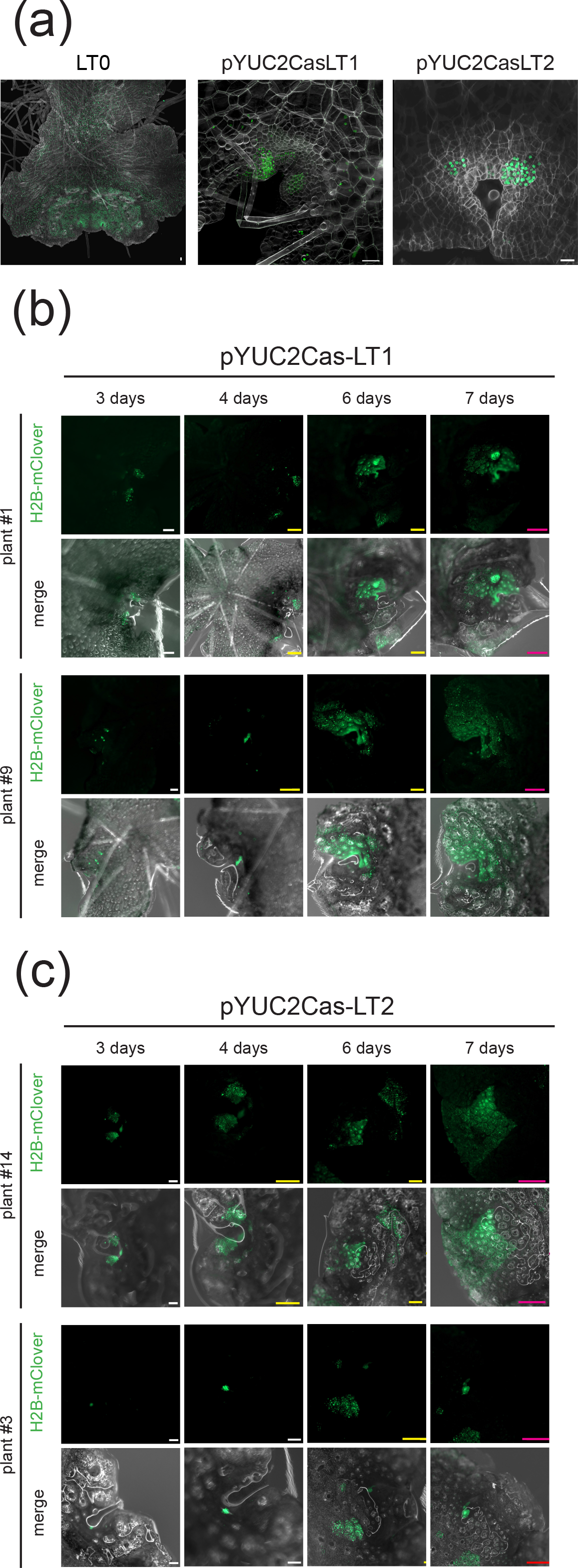
Functional test of the reporter lines in *Marchantia polymorpha*. Examples of fluorescent sectors in *M. polymorpha* expressing the lineage tracing reporters. (a) Confocal imaging of 3 days-old gemmae carrying the in-frame construct LT0 (left), the frameshift reporters LT1 (middle) or LT2 (right) with spCas9 under the control of promoter *pMpYUC2* active in the meristems of the apical notch. Bar = 20 μm. (b, c) Images from subsequent days after transferring gemmae from lines with LT1 (b) and LT2 (c) as in (a) onto fresh medium. # = individual transformants. White bar = 100 μm, yellow bar = 250 μm, magenta bar = 500 μm.

## DISCUSSION

The lineage tracing system based on frameshift correction after a targeted double strand break by Cas9 has proven its functionality *in planta*: it results in stable fluorescence labelling of nuclei in the original cell and all cells deriving from it, in different tissues of two different model plants. Depending on the time in development when the cut and repair happens, it labels single cells, short or longer cell files, or large sectors. These recapitulate and visualize known cell lineages, thereby offering confidence that the tool can help to systematically characterize also previously unknown lineages and their origin. The modular cloning system allows easy exchange of promoters for adaptation to other cell types and plant species. Moreover, the estradiol-inducible system adds a precise time control over initial frameshift corrections.

The organization of alternating vertical cell files in the Arabidopsis root epidermis, easily accessible to microscopic analysis, provided an extensively analysed lineage example and allowed comparison of our system with previously generated data. As the readout of this labelling is a lasting mark, it can convert a transient or very low level of transcription into a “yes or no” signal, for which frequencies can be quantified. We believe that lineage tracing is very suitable for producing a comprehensive visual map of differentiating cell files and tissue fate and help to characterize the role of key genes at developmental branch points.

Despite all advantages, there are several aspects that need to be considered when applying the lineage tracing reporters in cell types that might contribute to the development of the germline. The frequency of double strand cuts, frameshift-correcting and other mutations are important parameters. Once the target site is cut and repaired, regardless of whether with or without restoration of the frame, the reporter loses responsiveness, and too frequent events in the germline to the next generation might result in loss of usable lineage tracing lines. However, this risk appears modest as we observed a low frequency of germline events leading to functional H2B-mClover or complete loss of responsiveness, so that either genotyping of an amplicon across the target site or the easy and quick inspection of a few progeny plants can be used to exclude lines that are either fully labelled or do not have any sectors from somatic events. On the other hand, the accumulation of CRISRP/Cas9-induced mutations in the target site of the reporter offers the advantage to combine the visualization-based lineage tracing with a sequence-based system, using the created mutations as unique cell “barcode” for lineage reconstruction at single cell level. This increases the potential of our tool for comprehensive developmental studies even further.

While the reproducible occurrence of frameshift corrections is a strength of the system, too many events in somatic lineages can also be problematic. There might be applications when multiple events within the tissue can limit the spatial resolution of tracing single lineages. In these cases, the reporters could be modified by requiring alternative, less frequent repair pathways for the frameshift correction, e.g., alternative NHEJ which can result in fluorescence only through microhomology repair of the DSB. Again, the modular nature of the constructs allows for an easy exchange of the corresponding cassette.

Higher resolution and refinements are further possible by asking for the overlap between cell-specific expression of Cas9 and sgRNA from different PolII promoters. This is expected to label nuclei only in cells that contain both components at the same time, expanding the potential for tissue-specificity of the initial labelling. Future improvement and extension of the system could further include overlapping expression of multiple reporters with unique colour markers and unique sgRNA target sites. The reporter could also be designed to induce functional gain and loss of two FP reporters in the same transcript, to distinguish mono-from di-allelic events. The estradiol-inducible system works well and adds the option for precise timing of the labelling, and this can be combined with requiring additional specificity for the cooperation of all components. In addition to lineage tracing, the lines are also suitable reporters to study mutants in NHEJ repair pathways or to ask for conditions that influence the repair capacity. Another exciting application could be determining the mobility of RNAs encoding the CRISPR components in grafting experiments (Yang *et al.* 2023) by expressing the LT reporters in the scion and Cas9/gRNAs in the stock.

While the system was designed to address special developmental questions in plants, the principle of the lineage tracing system described here can be easily modified for application in other multicellular organisms. Given the rapid evolution of single-cell omics technology, the system described here provides a complementary approach to integrate simple visual readout and molecular tools to allow unprecedented resolution in functional description of multicellular development a single-lineage single-cell approach.

## EXPERIMENTAL PROCEDURES

### Construction of the reporter lines

All the binary vectors used in the present study were generated using the GreenGate modular cloning protocol (Lampropoulos *et al.* 2013) (Supplemental Table 1). To engineer a GreenGate-compatible destination vector, pSUN (Thomson *et al.* 2011) was linearized and the sulfadiazine resistance cassette replaced with the *ccdb* expression cassette, for positive selection of fully assembled vectors (Bernard 1996), flanked by GreenGate-compatible ends using Gibson Assembly^®^ Master Mix – Assembly (NEB, E2611). The resulting binary vectors pGGSun, despite a compact size of 6 kb, does not require the presence of helper plasmid pSoup and offers a high copy alternative to pGreenII-based destination vectors (Hellens *et al.* 2000).

The novel entry modules created for this study were cloned by insertion of PCR-amplified sequences in pJet1.3 or pMiniT 2.0 following manufacturer’s instructions (CloneJET PCR Cloning Kit, Thermo Scientific; NEB^®^ PCR Cloning Kit, NEB). *BsaI* recognition sites followed by module-specific overhangs were added to each primer used to amplify entry sequences. In case internal *BsaI* sites were present, they were removed using scar-free *BsaI*-cloning designed to introduce synonymous nucleotide substitutions (Lampropoulos *et al.* 2013).

The sgRNA expression cassette module was generated as tRNA-multiplex system described in (Bente *et al.* 2020) under the control of the U6-26 promoter. Briefly, all PCR-amplified component were assembled using *BsaI* cloning. The product was run on a 1.5% agarose gel in TAE buffer, the band with the expected size of a fully assembled fragment was purified (Zymoclean Gel DNA Recovery Kit, Zymo Research) and cloned in pJet1.3 as described above. The insert was then amplified with primers containing the module-specific overhangs and cloned as entry vector.

The Lineage Tracing reporter cassettes (LT0/LT1/LT2) were generated by cloning the H2B∷mClover∷UBQ10t sequence from pUBQ10∷H2B-mClover∷UBQ10t (Incarbone *et al.* 2022). The AAVS1 target sequence (Wang *et al.* 2015) and additional base pairs to modify the reading frame were introduced in the primer sequence. The final expression cassette was produced by adding either pHTR5 or pMpEF1a as promoter, for expression in *Arabidopsis thaliana* or *Marchantia polymorpha* respectively, using *BsaI* scar-free cloning.

The final assembly reactions were performed as follows. One μl of each module and destination plasmid were mixed with 1.5 μl of T4 DNA ligase buffer 10x, 1 μL T4 DNA ligase (400 U/μl) and 1 μl BsaI-HFv2 (M0202S, R3733S, NEB) and water to 15 μl. Reactions were incubated at 37°C for 5 min and 16°C for 5 min for 50 cycles, followed by 5 min at 50°C and 5 min at 80°C. Up to 5 μl of assembly reactions were used for heat-shock transformation of Stellar™ Competent Cells (Takara Bio).

All plasmids were purified with MiniPrep (Zyppy Plasmid Miniprep Kit, Zymo Research), and all vector sequences were confirmed with Sanger sequencing.

### Generation of transgenic plants

*Arabidopsis thaliana* accession Col-0 plants were grown in growth chambers with 16 h light/8 h dark cycles at 21°C/16°C and 130 μmol/m^2^s light intensity for approximately 4 weeks until they reached the flowering stage. They were then transformed via the floral dip method (Clough and Bent 1998) using *Agrobacterium tumefaciens* GV3101 pMP90 strain (Koncz and Schell 1986).

### *Marchantia polymorpha* strains and culture

*M. polymorpha* laboratory accessions Takaragaike-1 (Tak-1) and Takaragaike-2 (Tak-2) were used (Ishizaki *et al.* 2008). Sporangia were harvested and sterilized with 1% sodium dichloroisocyanurate for 4 min and then washed with sterile dH_2_O. Sporelings were transformed using the protocol described in (Ishizaki *et al.* 2008). Gemmae – vegetative propagules – were grown on solid ½ Gamborg medium in a growth chamber at 23°C under 24-hour 10-30 μmol m^−2^ s^−2^ white light.

### Crossings and analysis of plant material

To combine the lineage tracing reporter with the stem cell marker, plants bearing either pCLV3Cas-LT0, pCLV3Cas-LT1, or pCLV3Cas-LT2 constructs were crossed with plants carrying the pCLV3:H2B-mCherry reporter (Gutzat *et al.* 2020) to generate F1 hybrid seedlings.

Prior to imaging and induction experiments, seeds were aliquoted in open 2 ml tubes and surface-sterilized by 10 min exposure to chlorine gas produced by mixing 100 ml of 14% sodium hypochlorite with 10 ml of 37% HCl in a desiccator. Seeds were then plated either on agar-solidified germination medium (GM) or on soil and stratified at 4°C for two days to optimize germination. Plants were grown at 21°C with 130 μmol/m^2^s light intensity under either long day (16 h light/8 h dark) or short day (8 h light/16 h dark) conditions. To induce expression of spCas9 in plants carrying the inducible lineage tracing reporters, 7 days-old seedlings, grown on vertically positioned GM agar plates, were transferred to 35 mm diameter petri dishes containing 3 ml of liquid GM supplemented with 10 μM estradiol (β-Estradiol, E8875, Sigma Aldrich, freshly prepared by diluting a 1 mM stock solution in DMSO) or DMSO as mock treatment. After incubation, seedlings were washed in liquid GM and moved to either GM agar plates or microscopy slides with an imaging chamber (μ-Slide 2 Well Glass Bottom, cat 80287, Ibidi GmbH) and covered with a slice of solid medium.

### Microscopy and image processing

For the confocal imaging of Arabidopsis shoot apical meristems, plants at different developmental stages were dissected to isolate the apices. Twenty-five apices for each timepoint were then submerged in 700 μl fixing solution (2% methanol-free formaldehyde, 1 × MTSB, 0.1% Triton X-100) and placed under vacuum for 10 min at RT. Fixation was continued by incubating the samples at 37°C for 45 min. Samples were washed twice with 1x MTSB to remove excess fixative.

Fixed samples were then immersed in 1 ml of ClearSee solution (Kurihara *et al.* 2015), prepared by mixing 10% (w/v) xylitol, 15% (w/v) sodium deoxycholate and 25% urea (w/v) in distilled water, and stored at 4°C for at least 3 days. To ensure optimal clearing, the ClearSee solution was exchanged daily. For staining of cell walls, samples were submerged in ClearSee containing 0.1% (v/v) of SR2200 Cell Wall Stain (Renaissance Chemicals, UK) and incubated overnight in the dark at room temperature. Staining solution was discarded, and after two washes with fresh ClearSee, samples were mounted on microscopy slides in 200 μl of fresh ClearSee. Coverslips were added and sealed with clear nail polish and the slides stored at 4°C until imaging. To avoid damaging the meristem, a small chamber was created by placing vaseline drops on each corner of the coverslip facing the slide and gently pressed until the mounting medium was evenly distributed.

Images of plant samples were acquired using an inverted confocal microscope (Zeiss LSM980 or LSM 880 Axio Observer, Zeiss, Germany) equipped with an Airyscan detector. The images were taken using a 20x/0.8 plan-apochromat air objective coupled with a BP420-480+BP495-550 filter for SR2200 and mClover and BP420-480+BP570-630 filter for mCherry. The microscope was controlled with Zen Blue software (Zeiss, Germany). The images obtained were then processed with Zen Blue software for deconvolution and FIJI or Imaris (Oxford Instruments) software for orthogonal projection.

For time-lapse experiments of estradiol-induced lineage tracing, seedlings were mounted as described above. Images of the whole root were acquired every hour for a total of 20 h using Zeiss LSM980 inverted confocal microscope with an Airyscan II detector and controlled by Zen blue software.

Images of live plants were taken with a Lumar fluorescence stereomicroscope equipped with HPX120 (Zeiss, Germany).

## SUPPORTING INFORMATION

### Supplemental Figure 1

(a) Images of gemmae with the LT0 vector or non-transgenic gemmae (wt) at different time points. Bar = 100 μm

(b, c) 3D reconstruction and orthogonal view from confocal imaging of 3-days-old gemmae carrying the lineage tracing vectors pYUC2Cas-LT1 (b) and pYUC2Cas-LT2 (c). Bar = 30 μm

### Supplemental Table 1

List of binary and entry vectors used to generate the plasmids with the GreenGate system.

### Supplemental Table 2

List of primers used in the present study.

## ACKNOWLEDGEMENTS

The authors are grateful to Marco Incarbone for cloning components and to Zsuzsanna Mérai for useful comments on the manuscript. We appreciate the excellent support from the BioOptics Facilities at the Vienna BioCenter, especially Pawel Pasierbeck and Alberto Moreno Cencerrado. We also thank the Plant Facilities for good care of the plants, the Molecular Biology Services for reagents, and the Health Care Unit for making lab work possible even during the pandemic. The project was financially supported by the Vienna Science and Technology Fund (WWTF LS13-057).

## DATA AVAILABILITY STATEMENT

The manuscript contains all data and does not need a reference to any deposited data.

