## Supplementary figures and images for "A versatile CRISPR-based system for lineage tracing in living plants"

### Supplemental Figure 1

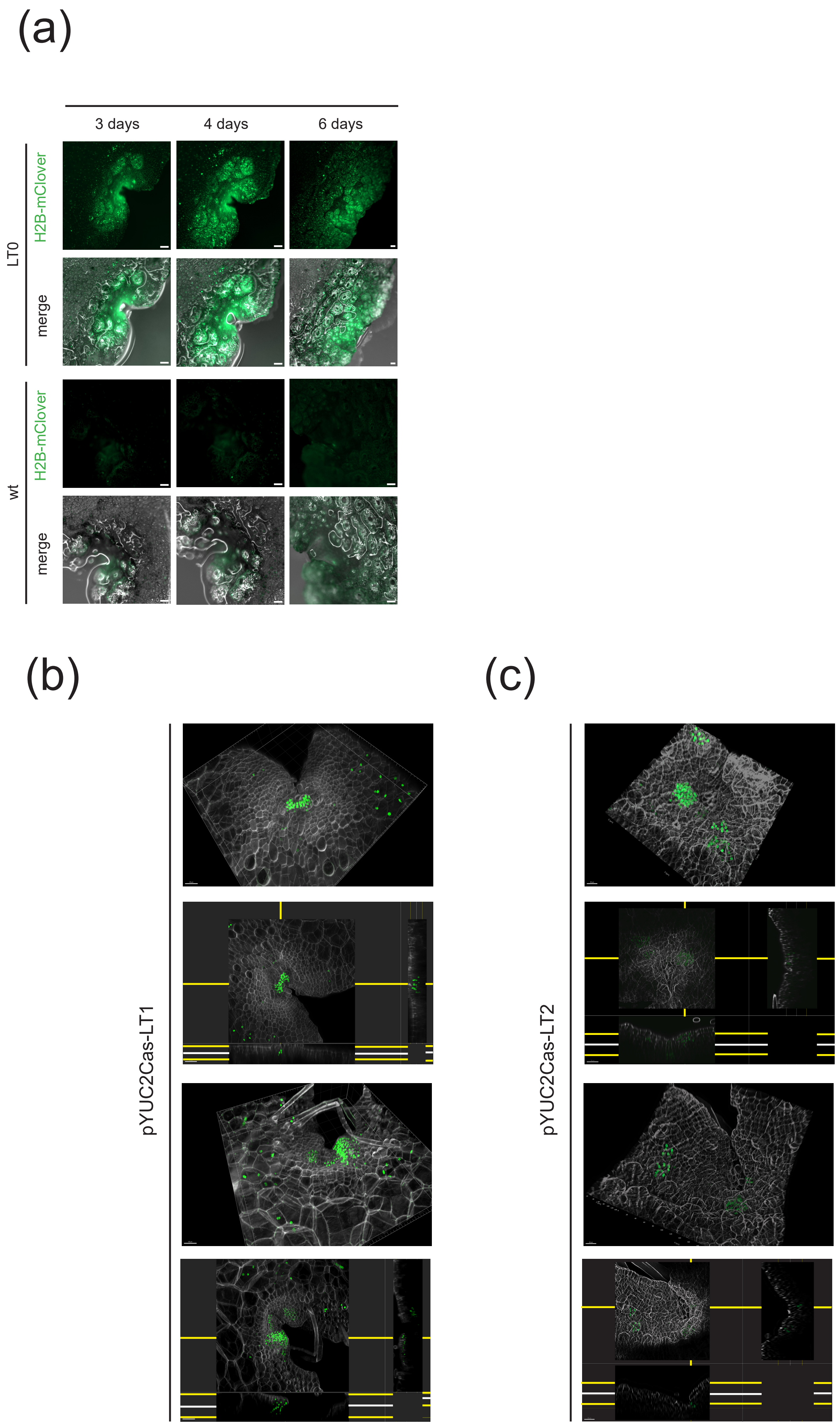


Supplemental Figure 1
